# Increase in egg resistance to desiccation in springtails correlates with blastodermal cuticle formation: eco-evolutionary implications for insect terrestrialization

**DOI:** 10.1101/767947

**Authors:** Helena Carolina Martins Vargas, Kristen A. Panfilio, Dick Roelofs, Gustavo Lazzaro Rezende

## Abstract

Land colonization was a major event in the history of life. Among animals, insects exerted a staggering terrestrialization success, due to traits usually associated with post-embryonic life stages, while the egg stage has been largely overlooked in comparative studies. In many insects, after blastoderm differentiation, the extraembryonic serosal tissue wraps the embryo and synthesizes the serosal cuticle, an extracellular matrix that lies beneath the eggshell and protects the egg against water loss. In contrast, in non-insect hexapods such as springtails (Collembola) the early blastodermal cells synthesize a blastodermal cuticle. Here, we investigate the relationship between blastodermal cuticle formation and egg resistance to desiccation in the springtails *Orchesella cincta* and *Folsomia candida*, two species with different oviposition environments and developmental rates. The blastodermal cuticle becomes externally visible in *O. cincta* and *F. candida* at 22 and 29% of embryogenesis, respectively. To contextualize, we describe the stages of springtail embryogenesis, exemplified by *F. candida*. Our physiological assays then showed that blastodermal cuticle formation coincides with an increase in egg viability in a dry environment, significantly contributing to hatching success. However, protection differs between species: while *O. cincta* eggs survive at least 2 hours outside a humid environment, the survival period recorded for *F. candida* eggs is only 15 minutes, which correlates with this species’ requirement for humid microhabitats. We suggest that the formation of this cuticle protects the eggs, constituting an ancestral trait among hexapods that predated and facilitated the process of terrestrialization that occurred during insect evolution.

**Research Highlights:**

1. The formation of the blastodermal cuticle produced during early embryogenesis coincides with a higher protection against water loss in springtail (Collembola) eggs.
2. *Orchesella cincta* eggs are more resistant to drought than *Folsomia candida* ones.
3. The formation of a protective egg cuticle would be an ancestral trait among hexapods that facilitated their process of terrestrialization.

**Graphical Abstract:** 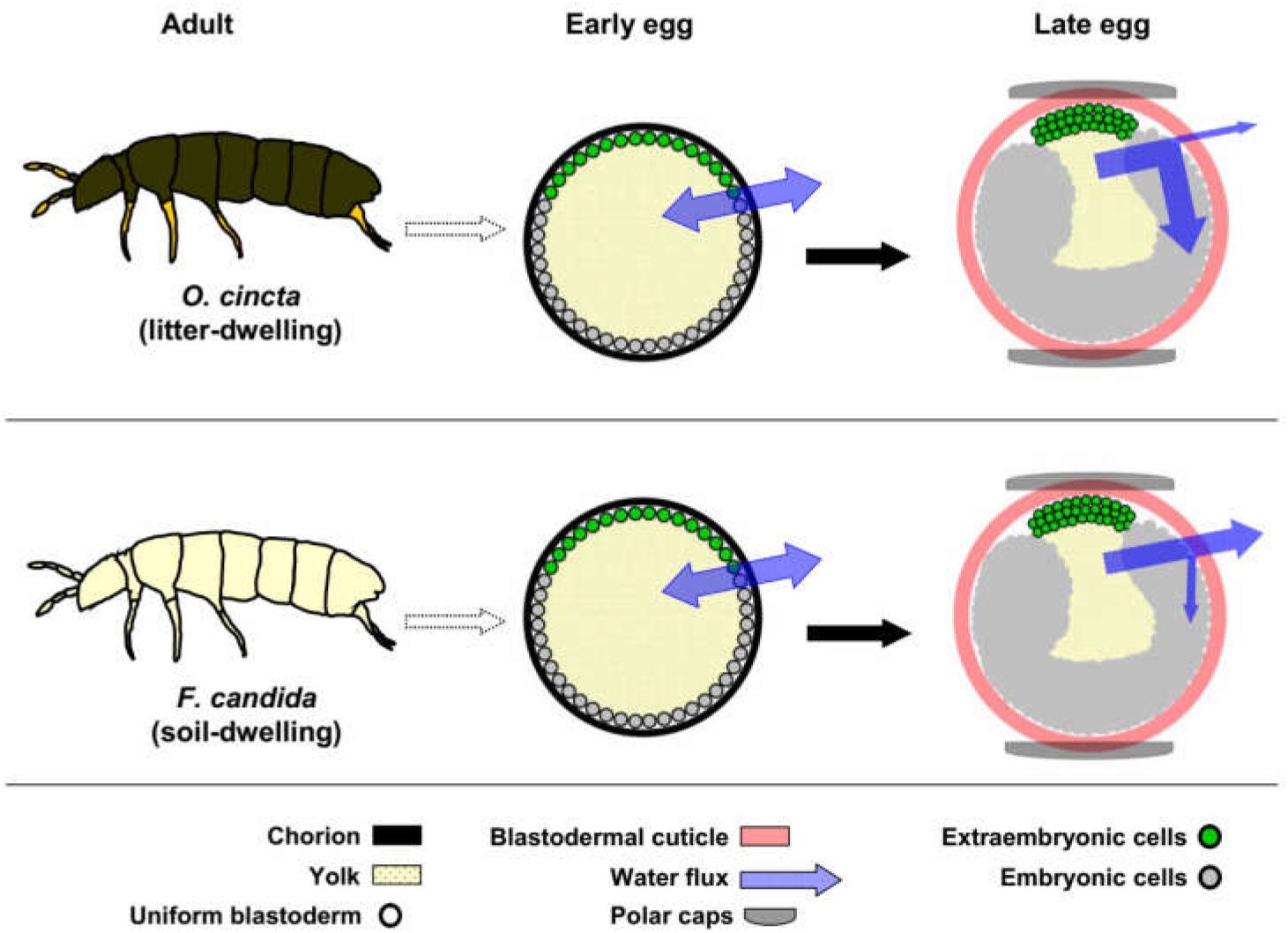

**Graphical Abstract legend:** Eggs when laid uptake water but are also prone to water loss. Late eggs acquire some protection against water loss, but at different levels, depending on the species.

## Introduction

Life originated in a marine environment and subsequent colonization of the land was a significant event for the evolution of many organisms on Earth, such as arthropods (Little, 1983; Selden, 2016). The arthropod phylum is comprised of the paraphyletic “Crustacea” and the monophyletic Chelicerata, Myriapoda and Hexapoda (constituted of Collembola, Protura, Diplura and Insecta) (Kukalová-Peck, 1987; Misof et al., 2014) (Suppl. Figure 1). It is estimated that multiple, independent terrestrialization events occurred during arthropod evolution, standing out the ones of hexapods, myriapods and chelicerates (Dunlop et al., 2013; Grimaldi and Engel, 2005; Rota-Stabelli et al., 2013). Specifically in the case of insect evolution, colonization of land led to an unparalleled diversification among animals. There are 1,527,660 described species of extant animals; arthropods and insects account for 79% and 66% of the total, respectively (Suppl. Figure 1). In other words, considering all living animal species, two-thirds are insects (Zhang, 2011).

In order to survive on land, insects evolved traits that allow them to overcome low or irregular water availability, such as an internalized respiratory system and an exoskeleton that minimizes water loss (Dunlop et al., 2013; Hadley, 1994; Little, 1983). Moreover, in addition to juvenile and adult characteristics, it has been suggested that traits related to the egg stage contributed to their successful terrestrialization (Zeh et al., 1989). Freshly laid insect eggs are covered by eggshell layers that are produced during oogenesis within the maternal body. These maternal eggshell layers will be collectively termed here as ‘chorion’. During the early stages of insect embryogenesis (Figure 1A-E), the blastoderm (Figure 1A) differentiates into the embryo proper and two extraembryonic tissues, serosa and amnion (Figure 1B). After differentiation, the serosal cells envelop the embryo (Figure 1C, D) and, in many species, secrete the serosal cuticle (Figure 1E) (Handel et al., 2000; Panfilio, 2008; Rezende et al., 2016, 2008). This cuticle is a stratified extracellular matrix structure composed of a thin external epicuticle and a thick internal endocuticle. It is assumed that the serosal epicuticle is composed of proteins and hydrophobic compounds (*e*.*g*. lipids, waxes, hydrocarbons) while the serosal endocuticle contains proteins and chitin (Jacobs et al., 2013; Rezende et al., 2016). This post-zygotic serosal cuticle constitutes the innermost eggshell layer (located below the chorion) and is fundamental to protect the egg against water loss (Jacobs et al., 2013; Rezende et al., 2016, 2008). The presence of two separate extraembryonic tissues (serosa and amnion) is an insect novelty (Wheeler, 1893) and together with the serosal cuticle are considered as crucial traits that fostered the evolution of insects on land (Jacobs et al., 2013; Zeh et al., 1989). Other arthropods have one extraembryonic tissue that never wraps the embryo (Anderson, 1973; Jacobs et al., 2013), yet terrestrial embryogenesis in these species faces the same environmental challenges, highlighting the need for comparative studies with other lineages.

**Figure 1:**
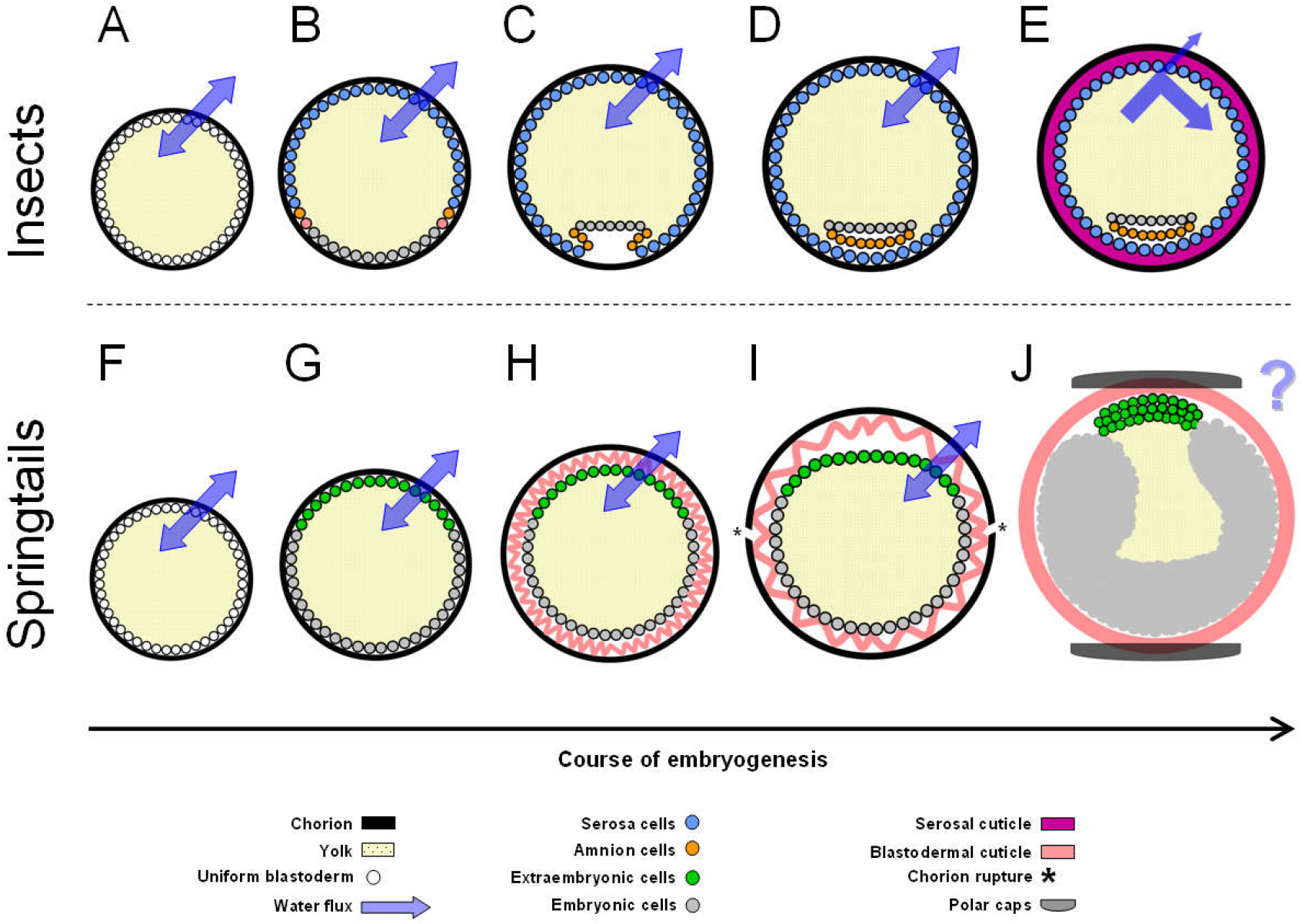
Dynamics of post-zygotic cuticle formation during early embryogenesis of insects and springtails. Schemes of egg cross-section. Increases in egg sizes are due to water uptake. (**A**) Insect egg at early embryogenesis with cells at the uniform blastoderm stage. (**B**) The blastoderm differentiates into embryonic cells and two extraembryonic tissues: serosa and amnion. (**C, D**) Serosal cells wraps the yolk, embryo and amnion. (**E**) The serosa produces its cuticle, which increases the egg protection against water loss. In this example the insect egg absorbs water, but this is not a universal trait. Based and reviewed on Rezende et al. (2016). (**F**) Springtail egg at early embryogenesis with cells at the uniform blastoderm stage. (**G**) Blastoderm cells differentiate into embryonic and extraembryonic ones. (**H**) The differentiated blastoderm produces a blastodermal cuticle that is initially wrinkled. (**I**) Extraembryonic cells move inward the egg while the increase in egg volume leads to chorion rupture. (**J**) Chorion remnants become the polar caps and the expanded blastodermal cuticle is now the sole egg cover. At this stage, egg permeability to water is unknown. For the sake of simplicity, yolk cells are not shown. Depending on the species, the blastodermal cuticle is secreted before or after blastoderm differentiation, is smooth or with ornaments, is one or more cuticles. The degree of egg size increase varies among species, as well as the embryonic morphological stage where chorion rupture occurs. Based on Bretfeld (1963), Jura (1972), Jura et al. (1987), Krzysztofowicz (1986a), Krzysztofowicz (1986b), Marshall and Kevan (1962), Prymus-Naczynksa (1978), Prowazek (1900), Tomizuka and Machida (2015), Uemiya and Ando (1987b), Uemiya and Ando (1987a). Since the serosa and the amnion are insect novelties (see Introduction), the non-embryonic cells of the springtail blastoderm are simply named as “extraembryonic” (Anderson, 1973; Jacobs et al., 2013). These extraembryonic cells will become the dorsal organ (Jura, 1972) but, for the sake of clarity, this process is not included here.

In fact, the production of supplementary eggshell layers during embryogenesis is documented for species within all other arthropod groups where these eggshell layers are referred to as the blastodermal cuticle (also termed blastodermic cuticle or blastoderm cuticle) (Machida, 2006; Rezende et al., 2016). For this trait, the Collembola offer an excellent research model as a hexapod group closely related to the insects but lacking such a tremendous diversification: collembolans, colloquially called ‘springtails’, account for only 0.5% of the total species number of living animals (Suppl. Figure 1). Moreover, collembolan egg development has been well described for many species in classical, morphological studies on embryogenesis, eggshell structure, and specifically blastodermal cuticle formation (Bretfeld, 1963; Jura, 1972; Jura et al., 1987; Krzysztofowicz, 1986a, 1986b; Larink and Biliński, 1989; Marshall and Kevan, 1962; Prowazek, 1900; Prymus-Naczyńska, 1978; Tamarelle, 1981; Tiegs, 1942; Tomizuka and Machida, 2015; Uemiya and Ando, 1987a, 1987b; Wojtowicz, 1978). Briefly, during early embryogenesis (Figure 1F-J) the springtail blastoderm (Figure 1F) differentiates into two types of cells: the extraembryonic tissue and the embryonic rudiment (Figure 1G). Depending on the species, cells of either the uniform undifferentiated blastoderm or the differentiated blastoderm secrete the blastodermal cuticle (Figure 1H). In parallel, water uptake increases egg volume, which leads to chorion rupture (Figure 1I), thus forming the polar caps (*i.e.* chorion remnants that lie at opposite poles of the egg). At this time the blastodermal cuticle becomes the sole protective egg layer (Figure 1J).

The extraembryonic cells will give rise to the dorsal organ (Jura, 1972; Tamarelle, 1981; Tiegs, 1942), a process that will not be detailed here. The number of blastodermal cuticles may vary from one to four layers, depending on the species (Jura, 1972). Although the blastodermal cuticle is generically described as a resistant barrier that protects the egg (Machida, 2006; Rezende et al., 2016; Tiegs, 1942) “*nothing is actually known as to their permeability to water or air*”, as stated by Jura (1972).

In any case, springtail blastodermal cuticles are structurally similar to serosal ones, at least for those species where data are available: a thin epicuticle layer and a thick stratified endocuticle layer composed of lamellae with alternating electron-dense and electron-lucent layers (Krzysztofowicz, 1986a, 1986b; Lamer and Dorn, 2001; Prymus-Naczyńska, 1978; Rezende et al., 2016; Tamarelle, 1981). We thus hypothesize that the blastodermal cuticle would also protect the springtail egg against water loss.

Here, we analyze this relationship through physiological correlation and desiccation assays in two springtails species, *Orchesella cincta* (Entomobryidae) and *Folsomia candida* (Isotomidae). Both species are used as biological indicators in soil ecotoxicology (Hopkin, 1997; Fountain and Hopkin, 2005) and are supported by genome sequence resources (Faddeeva-Vakhrusheva et al., 2016; Faddeeva-Vakhrusheva et al., 2017). Importantly, these species also differ in their habitat, with litter-dwelling (epidaphic) *O. cincta* experiencing a drier environment than the soil-dwelling (eudaphic) *F. candida*. In fact, the overall water management capacity of *Folsomia candida* eggs were recently described (Holmstrup, 2019). Water content, water uptake and volume increase throughout development were assessed. In addition, drought exposure led to water loss and decrease in viability. Eggs are more sensitive than juvenile and adult forms, with no hatching occurring at the wilting point of plants (98.9% relative humidity, corresponding to a water potential of −1.5 MPa). However, these data were not associated with stage-specific developmental changes occurring in the eggs, such as blastodermal cuticle formation and chorion rupture

To deepen our understanding of water relations in springtail eggs and foster the development of resources for ongoing work across research groups that focus on these species, we described in detail the rearing of these species and we investigated embryonic aspects related to the blastodermal cuticle formation and whether this structure could be involved in desiccation resistance. We found that blastodermal cuticle production coincides with an increase in egg protection against water loss in both species, although to distinct levels of protection.

## Methods

### Springtail life history, rearing conditions and resources

*Orchesella cincta* adults are pigmented and measure about 6 mm in length. Its embryogenesis lasts for about 6 days at 19-24 ºC, while egg size at the chorion rupture stage is about 200 - 250 μm in diameter, when two blastodermal cuticles are produced. This cuticle is covered with thorns (Bretfeld, 1963). Its postembryonic development takes at least 44 days before reaching the adult stage (Hopkin, 1997; Joosse et al., 1972). The species reproduces sexually through indirect sperm transfer via spermatophores. The individuals molt during adult life, and sperm cells cannot be stored by females. After each adult molt a female enters a new reproductive cycle and take up a new spermatophore (Hopkin, 1997).

*Folsomia candida* is a parthenogenetic species. Adults are unpigmented, blind and measure between 1 to 3 mm in length. Its embryogenesis lasts about 10 days and hatchlings take 24 days to reach the adult stage at 20 ºC. Its eggs are smaller than those of *O. cincta*: they measure about 80 μm when laid and increase in size up to 165 μm at the chorion rupture stage (Marshall and Kevan, 1962). Adults can survive under relatively dry conditions by increasing osmolality through production of sugars and polyols, facilitating absorbance of water vapor (Fountain and Hopkin, 2005; Hopkin, 1997; Roelofs et al., 2013).

Specimens of *O. cincta* and *F. candida* (Berlin strain) have been maintained for more than 20 years at the Department of Ecological Sciences, Animal Ecology Section, Vrije Universiteit (Amsterdam, The Netherlands) (Agamennone et al., 2015; Sterenborg and Roelofs, 2003). The *Orchesella cincta* population originates from several forests in The Netherlands (van Straalen et al., 1987). Whenever necessary, specimens from the wild were introduced in the laboratory culture to restore vigor. Both species were reared in a climatic room set at 15 ºC, 75% relative humidity (RH) and 16:8 hour light:dark photoperiod. *Orchesella cincta* was maintained in PVC jars (25 cm diameter) containing twigs and plaster of Paris (500g plaster of Paris and 500 ml demineralized water). Juveniles and adults were fed on algae grown over small pieces of twig kept in a humid environment (Suppl. Figure 2). *Folsomia candida* was kept in plastic jars (20 x 15 cm) filled with a mixture of plaster of Paris (Siniat-Prestia, multiple purpose plaster) and charcoal (CAS 7440-44-0, Merck, Germany) (500g plaster of Paris, 500 ml demineralized water and 50g charcoal) (Suppl. Figure 3). Juveniles and adults were fed with commercial yeast (Dr. Oetker, Bielefeld, Germany). Every two days, demineralized water was added into the jars of both species to keep the plaster moist.

### Temperature of the assays

Egg collection for the description of *F. candida* embryogenesis was undertaken at 21 ºC, which is the optimal hatching temperature (Fountain and Hopkin, 2005). All other assays were performed at 20 ºC at which *O. cincta* egg production is high (Joosse et al., 1972) and is the standard test temperature for *F. candida* in its use for soil toxicity and remediation applications (internationally standardized test ISO 11267; reviewed in Fountain and Hopkin, 2005).

### Egg laying procedures

Egg laying was performed according to the specific characteristics of each species’ reproduction process. For *O. cincta*, adults were collected from rearing jars, sexed and 10 couples were transferred to small circular jars (5 cm diameter) containing plaster of Paris or plaster of Paris-charcoal and a piece of bark peeled off of a twig (Suppl. Figure 4A). Adults were fed on algae grown on top of these bark pieces. Since a female starts a new reproductive cycle after each adult molt, the jars were checked daily to determine when they molted. After molting the jars were carefully observed to check the presence of eggs. As soon as the first eggs were observed, all animals were transferred to a new jar and egg laying occurred during eight or 12 hours. For each instance of egg collection, 10 small jars were used (a total of 100 couples). Plaster humidity was maintained by adding demineralized water every day.

For *F. candida*, large adult parthenogenetic females were collected from the rearing jars and transferred to plastic pots (10 x 8 cm) containing plaster of Paris-charcoal (Suppl. Figure 4B). Each pot contained 15-20 females and for each egg laying interval, 10 pots were used. All females were allowed to oviposit directly onto the plaster surface for eight hours. After this period, all females were removed from containers. Plaster humidity was maintained by adding demineralized water every two days. Egg laying for embryogenesis description was done differently (see below).

For egg laying intervals of eight or 12 hours, we report de median age, i.e. X±4 or X±6 hours old, respectively; X being the number of hours elapsed after the end of the egg laying period.

### Defining the total period of embryonic development

Groups of at least 20 eggs obtained as described above were left developing at 20 ºC and 75% RH and observed in the stereomicroscope at two and one hour intervals, for *O. cincta* and *F. candida*, respectively. For each species three independent experiments with at least three replicates each were performed. A total of 225 and 315 eggs were evaluated for *O. cincta* and *F. candida*, respectively. Eggs were maintained in moist conditions until the end of this experiment. The period of complete embryonic development was defined when 50% of the juveniles hatched.

### Analysis of chorion rupture

Replicates of 25 or 30 eggs were used to evaluate when chorion rupture occurs in both species. Every hour the number of eggs showing the exposed blastodermal cuticle and the polar cap (i.e. the ruptured chorion) was counted under a stereomicroscope. For *O. cincta* two independent assays were performed, one with two replicates, the other with three replicates with a total of 140 eggs employed. For *F. candida* four independent assays were performed, each with ate least three replicates, with a total of 365 eggs employed. Eggs were maintained in moist conditions until the end of this experiment. When 50% of the samples had their chorion ruptured was defined as the percentage of development at which this event occurs.

### Analysis of *F. candida* embryonic development

For this analysis, eggs were collected directly from the plaster-charcoal dishes in which the *F. candida* colony was maintained, without removing juveniles or adults. Eggs at different ages were collected with a small, pruned paintbrush manipulated down a dissecting microscope (Wild M5A Heerbrugg) and stored on damp filter paper in a culture dish until fixed. Collected eggs were transferred from the damp filter paper to a 1.5 ml tube of water with the paintbrush. The tube was centrifuged at 6000 rpm for 30 seconds, to limit the number of eggs floating at the surface. Eggs were then washed with PBT (PBS with 0.1% Tween-20) and subsequently treated with bleach (50% bleach in PBS) for 2.5 minutes. Embryos were washed three times with PBT and fixed for 25 minutes in a fixing solution (5% formaldehyde / PBT) on a shaker. Eggs were then washed once in PBT. To improve eggshell permeablization for staining, eggs were sonicated for 10 seconds with a probe sonicator (Misonix XL series, power 2-4). To visualize developmental morphology by nuclear staining, eggs were stained for 1 hour with 1 μg/ml DAPI (Sigma) diluted in PBT followed by 4 PBT washes of 15 minutes each, at room temperature. Eggs were mounted in 70% glycerol/PBS and imaged with DIC and fluorescence optics on an upright epifluorescent microscope (Zeiss Axiophot with a Hamamatsu color camera, model C4742-95-12NRB).

### Analysis of egg permeability to water

To test if water permeability changes in springtail eggs during embryogenesis, eggs samples at different stages of development were analyzed. For *O. cinta*, groups of 20 or 25 eggs were collected from moist plaster with a paint brush and placed onto a polycarbonate filter (25 mm diameter, 8 mm pore, Poretics Corporation), deposited on a drop of demineralized water. The polycarbonate filter with eggs was blotted onto a dry filter paper and left air-drying for two periods: 15 minutes or 2 hours. Temperature and relative humidity of the assays varied between 21.2 - 23.9 ºC and 45 - 76%, respectively. It was not possible to control the relative humidity of the room, but we observed that humidity variation did not affect the outcome of this experiment. For each stage of embryogenesis the number of intact or shriveled eggs was counted under a stereomicroscope. A similar procedure was performed with *F. candida*; however groups of 15 or 20 eggs were left air-drying for 5 or 15 minutes at 22 ºC and 46 - 58% of relative humidity. A total of 890 and 1,435 eggs were evaluated for *O. cincta* and *F. candida*, respectively. When 50% of the eggs are protected against shriveling was defined as percentage of development at which this event occurs.

### Analysis of egg viability under dry conditions

Group of eggs, at different stages of embryogenesis, were transferred from moist plaster onto a polycarbonate filter (25 mm diameter, 8 mm pore, Poretics Corporation), blotted onto a dry filter paper and left air-drying for 15 minutes or 2 hours (see Suppl. Figure 5). After these periods, eggs were returned to moist plaster until embryogenesis completion, when egg viability was checked through juvenile hatching. For each species and each stage of embryogenesis at least two independent assays were performed, each with at least two replicates. Each replicate consisted of at least 15 and 25 eggs for *O. cincta* and *F. candida*, respectively. A total of 960 and 1,655 eggs were evaluated for *O. cincta* and *F. candida*, respectively. Control replicates kept in moist plaster throughout embryogenesis were used to normalize juvenile hatching, summing up a total of 585 and 755 eggs of *O. cincta* and *F. candida*, respectively. Temperature and relative humidity varied between 21.2 - 23.9 ºC and 45 - 78%, respectively. It was not possible to control the relative humidity of the room, but we observed that humidity variation did not affect the outcome of this experiment.

### Statistical analyses

For all experiments, mean and standard error were calculated. One-way analysis of variance (ANOVA) (P < 0.0001) followed by Tukey’s Multiple Comparison Test was performed in the assay of egg viability under dry conditions.

## Results

### Egg laying behavior and total time of embryonic development differs between springtail species

While While both springtail species have been described in previous literature, here we report on general features for obtaining eggs and conducting physiological experiments under laboratory-controlled conditions (see Methodology and Suppl. Figs. 2-5). We observed consistent species-specific characteristics in oviposition behavior and general fecundity. Eggs of *O. cincta* are laid individually or in pairs, on the top of the bark or within its indentations (Figure 2A). In contrast, the eggs of *F. candida* are laid in clutches, preferentially in small crevices in the plaster substrate (Figure 2B, Suppl. Figure 3A, B), consistent with this species’ eudaphic life style. Due to a higher rate of egg production in *F. candida*, we were able to analyze particularly large sample sizes of this species in each assay.

**Figure 2.**
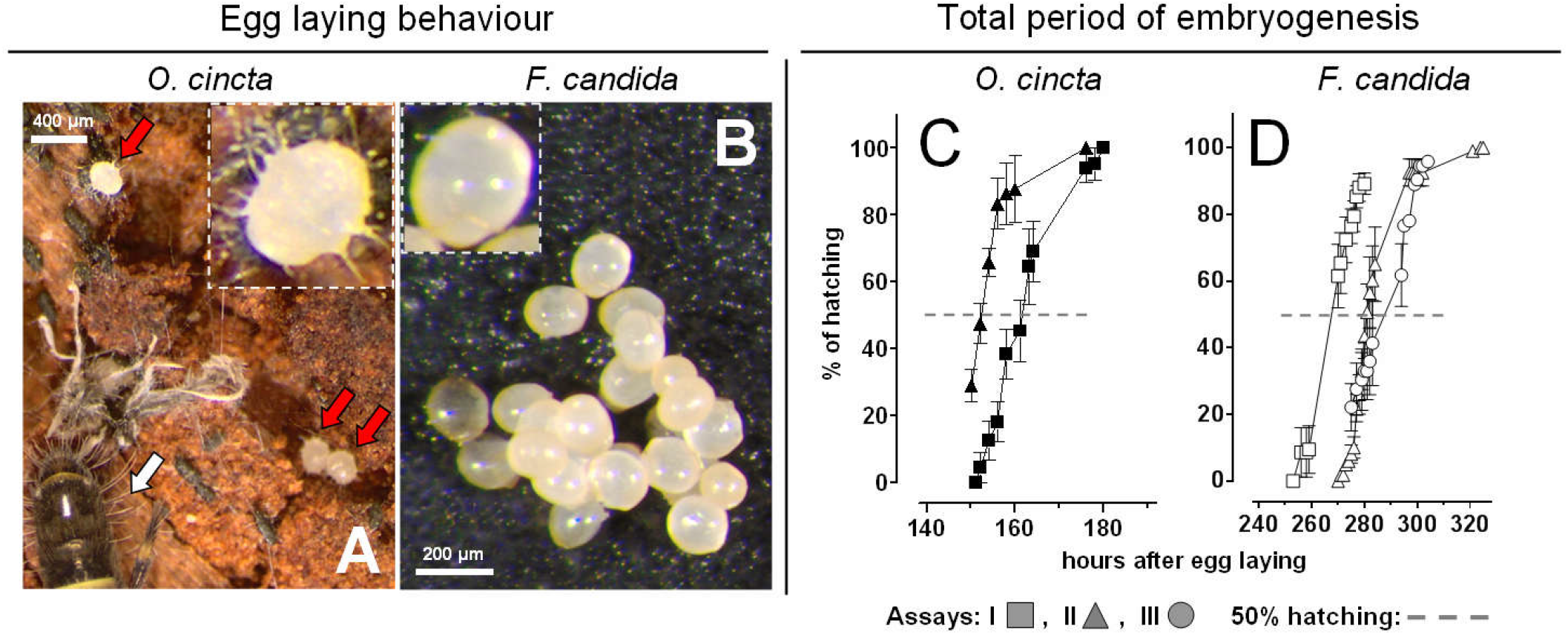
Egg laying behavior and total period of embryogenesis differs between species. (**A**) Eggs of *O. cincta* (red arrows) are laid singly or in pairs. Chorion rupture already occurred in these eggs. White arrow: posterior part of an adult body. (**B**) Eggs of *F. candida* are laid in a cluster. Insets show magnified eggs. As described in the Introduction, *O. cincta* eggs are larger than *F. candida* ones. (**C**, **D**) Cumulative juvenile hatching is depicted for eggs developed at 20 ºC on moist plaster. Embryogenesis occurs faster in *O. cincta* (**C**) than in *F. candida* (**D**). For each independent assay, plot points represent the mean and standard error, normalized by total hatching. A total of 225 and 315 eggs of *O. cincta* and *F. candida* were evaluated, respectively.

To precisely establish the quantitative framework for our physiological assays on blastodermal cuticle function, we next determined the duration of embryogenesis in each species, from oviposition to hatching (Figure 2C, D). *Orchesella cincta* has nearly twice as fast a developmental rate as *F. candida*: *O. cincta* juveniles hatch at 156 ± 5 hours after egg laying (HAE) (approximately 6.5 days), with 94 ± 4% viability whereas *F. candida* juveniles hatch at 278 ± 11 HAE (approximately 11.6 days), with 90 ± 10% viability. Given the differences between the total periods of embryogenesis, subsequent results are presented in percentages of embryogenesis to allow for direct comparison between species

### Maternal chorion rupture occurs after the first quarter of embryogenesis

The process of blastodermal cuticle formation during collembolan embryogenesis and the subsequent rupture of the maternally laid chorion have been characterized morphologically by light and electron microscopy (see Introduction and Figure 1). Here, we investigated the timing of chorion rupture during embryogenesis as an external morphological indication of when this cuticle has formed and expanded (Fig. 3).

**Figure 3.**
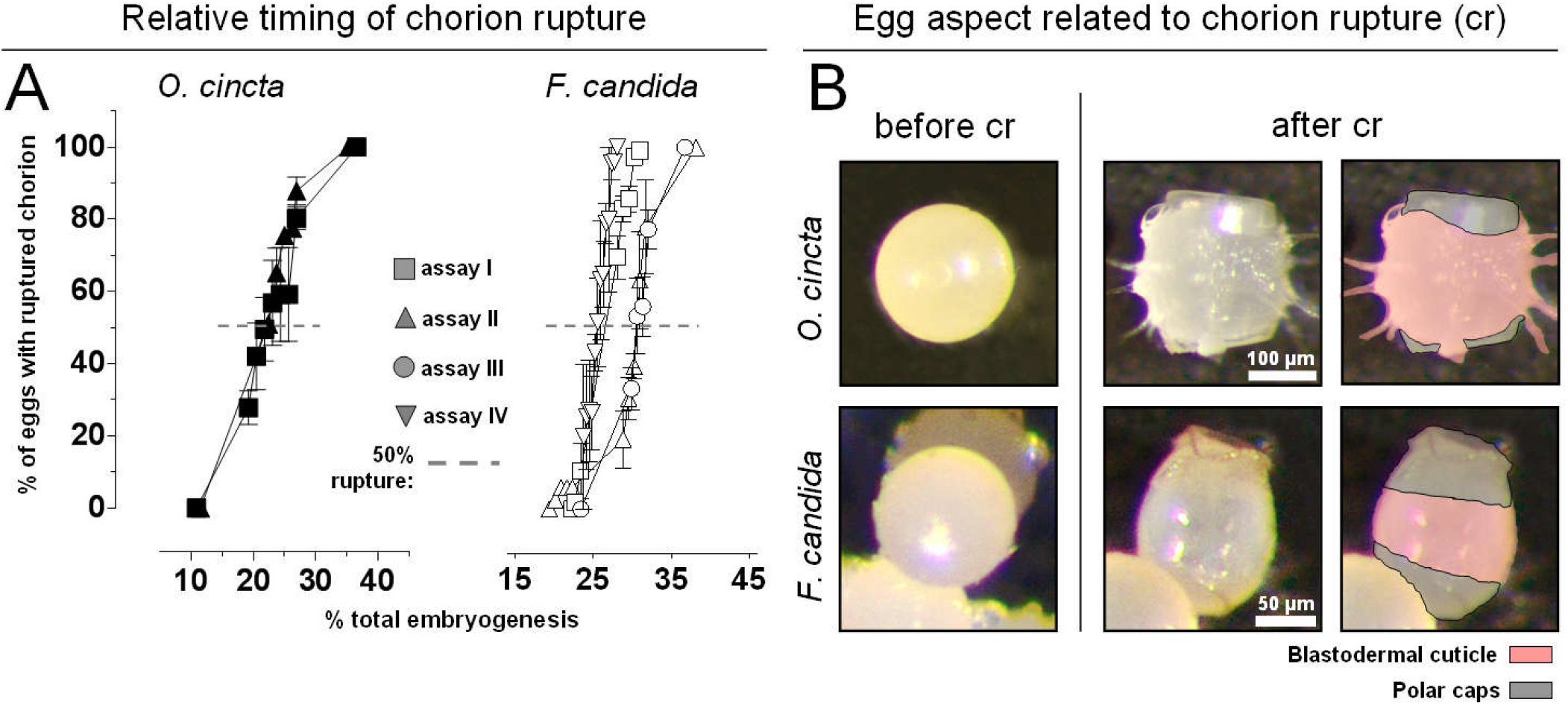
Chorion ruptures and blastodermal cuticle is exposed within the first quarter of embryogenesis. For both species eggs developed at 20 ºC. (**A**) Chorion rupture of *O. cincta* and *F. candida* eggs were recorded. Each geometric shape represents an independent assay and each point represent mean and standard error, normalized by total eggs employed per assay. A total of 140 and 365 eggs of *O. cincta* and *F. candida* were evaluated, respectively. (**B**) Egg aspect before and after chorion rupture. Pseudocolour in right-most panels indicate the polar caps (grey) and the exposed blastodermal cuticles (in pink)

For both species chorion rupture occurs after the first quarter of embryogenesis (Figure 3A). In *O. cincta* the chorion ruptures at 22% (~ 35 HAE) and in *F. candida* at 28% (~ 78 HAE) of total embryonic development. This period was defined as when 50% of all chorions ruptured. Before chorion rupture, eggs of both species are spherical and with a smooth surface (Figure 3B, left panels). After chorion rupture the blastodermal cuticle of *O. cincta* has thorns while the *F. candida* one is smooth (Figure 3B, right panels, Suppl. Figure 3C). *O. cincta* blastodermal cuticle is more rigid: after juvenile hatching, the cuticle maintains its form and resembles an empty bowl, while in *F. candida* this cuticle is pliable, losing its shape after juvenile hatching.

### Blastodermal cuticle formation is the first major event in F. candida embryogenesis

While blastoderm cuticle formation is completed after a quarter of total embryogenesis has elapsed, by definition the blastodermal stage is early, immediately following fertilization and the first cleavages (Jura, 1972). For context, here we document *F. candida* embryonic development from the blastoderm stage until hatching (Figure 4). Cells of the undifferentiated blastoderm divide (Figure 4A, B) before giving rise to the germ band (*i.e.,* the embryo proper) and the extraembryonic tissue (Figure 4C, D). Chorion rupture occurs between the stages depicted in Figure 4B and 4C. *Folsomia candida* eggs have different mechanical properties for fixation and permeabilization (through sonication) before (Figure 4A-B) and after chorion rupture (Figure 4C-H), with the pre-rupture stages more sensitive to damage. The germ band is initially formed on the surface of the yolk with the ventral surface facing outward but then it sinks into the yolk, assuming a concave posture (compare Figure 4D and E’). As embryonic development progresses, morphological structures including the segments and appendages mature and become more defined while the remaining yolk is restricted to the dorsal part of the egg (Figure 4E-F’), where eventually it is taken up within the gut (Figure 4G, G’). The external form of the body is completed during epidermal dorsal closure (Figure 4H, H’). After hatching, the whole body of the eyeless juvenile expands (Figure 4I). The description of *F. candida* embryogenesis falls within the general pattern of springtail egg development previously described for other species (see Introduction).

**Figure 4:**
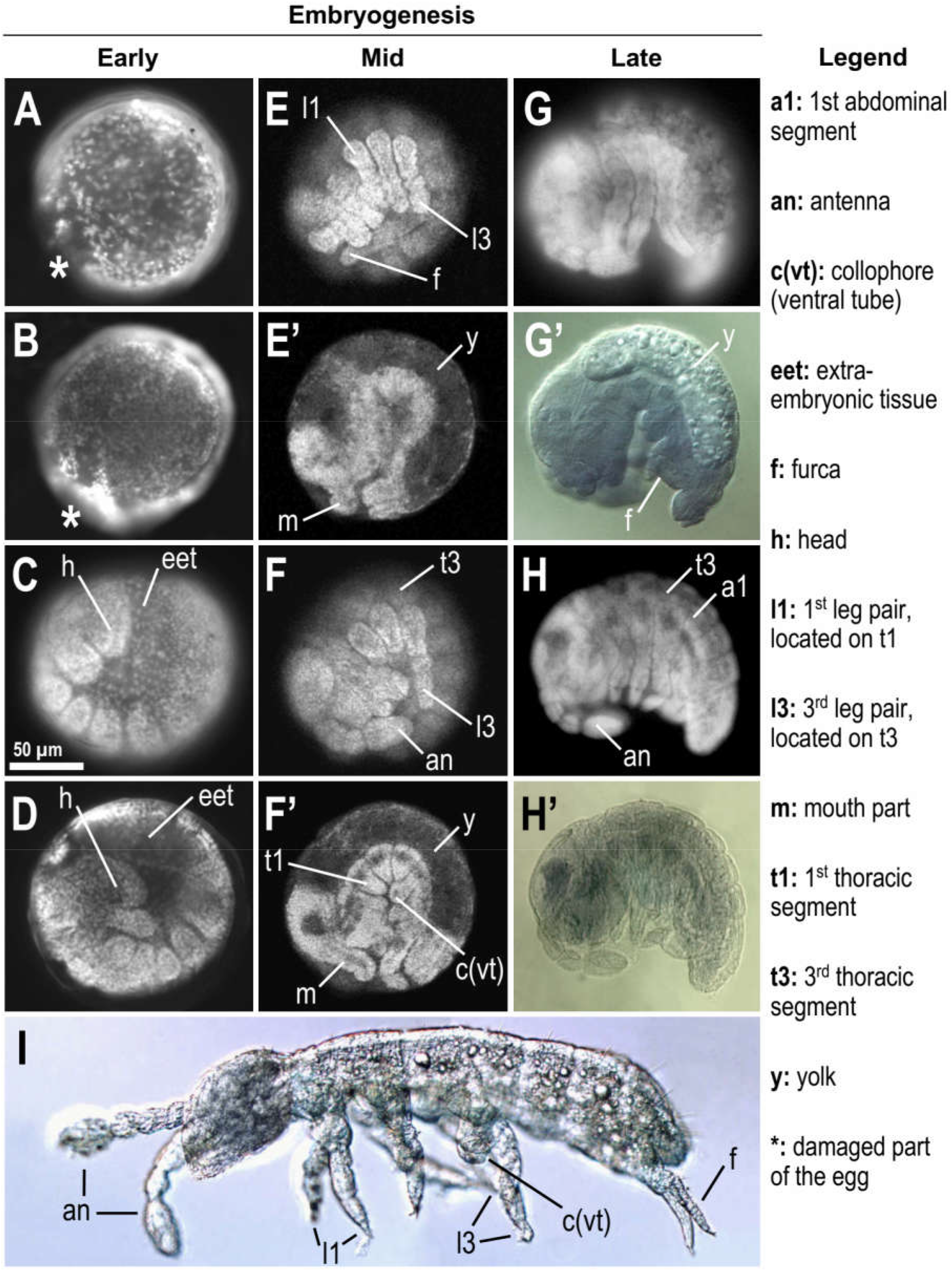
Stages of *Folsomia candida* embryogenesis. Images of all panels with the exception of G’, H’ and I: fluorescence microscopy, where cell nuclei were visualized with DAPI or propidium iodide. Images of panels G’, H’ and I: DIC microscopy. In all images anterior is to the left and dorsal is up. (**A**) Early undifferentiated blastoderm stage. (**B**) Late undifferentiated blastoderm stage. (**C**-**D**) Germ band at consecutive stages with increasing number of segments being defined. Between **D** and **E** the germ band sinks into the yolk shifting from a convex to a concave shape. (**E**-**E’**) Egg shown in two different focal planes; morphological definition of the mouth parts, legs and furca are evident. (**F**-**F’**) Egg shown in two different focal planes: antenna, ocular structure and the collophore (ventral tube) are evident. (**G**-**G’**) Egg at the stage after dorsal closure. In the focal plane in G’ a thin layer of epidermis is discernible as a complete dorsal cover over the granular yolk. (**H**-**H’**) Pharate juvenile at the end of embryogenesis. (**I**) Eyeless juvenile shortly after hatching. Indicative scale bar of 50 μm in panel C is based on egg size after chorion rupture (typically 150 μm; maximum diameter ~ 165 μm).

### Formation of the blastodermal cuticle coincides with an increase in the egg protection against water loss

Given the early commitment of springtail embryos to cuticle production prior to subsequent development, we next physiologically tested if the timing of cuticle formation is associated with change in the egg’s capacity to withstand water loss. To assess water retention, eggs were desiccated in air at different stages of embryogenesis (Figure 5). Eggs of *O. cincta* were exposed to dry conditions for 15 minutes or two hours (Figure 5A, B). Under both treatments, we observed a marked increase in egg protection after the blastodermal cuticle had formed. In contrast, *F. candida* eggs are less robust and all of them shriveled in the 15-minute assay (Figure 5D). However, when we reduced this to a five-minute assay, we could indeed observe an increase in water loss protection associated with the blastodermal cuticle (Figure 5C). Based on the stage when 50% of the eggs are intact (*i.e.* without shriveling) in Figure 5A and C, we assume that the blastodermal cuticle is formed in *O. cincta* at 15% (23 HAE) and in *F. candida* at 22% (~ 62 HAE) of embryogenesis. Consistent with the necessity to maintain a functional protective cover throughout embryogenesis, we therefore demonstrate that blastodermal cuticle formation is correlated to a protection against water loss before the chorion ruptures, with 6-7% development or half a day’s difference (0.5-0.7 d) between these two events (Table 1); compare also Figures 3A and 5A, C.

**Figure 5:**
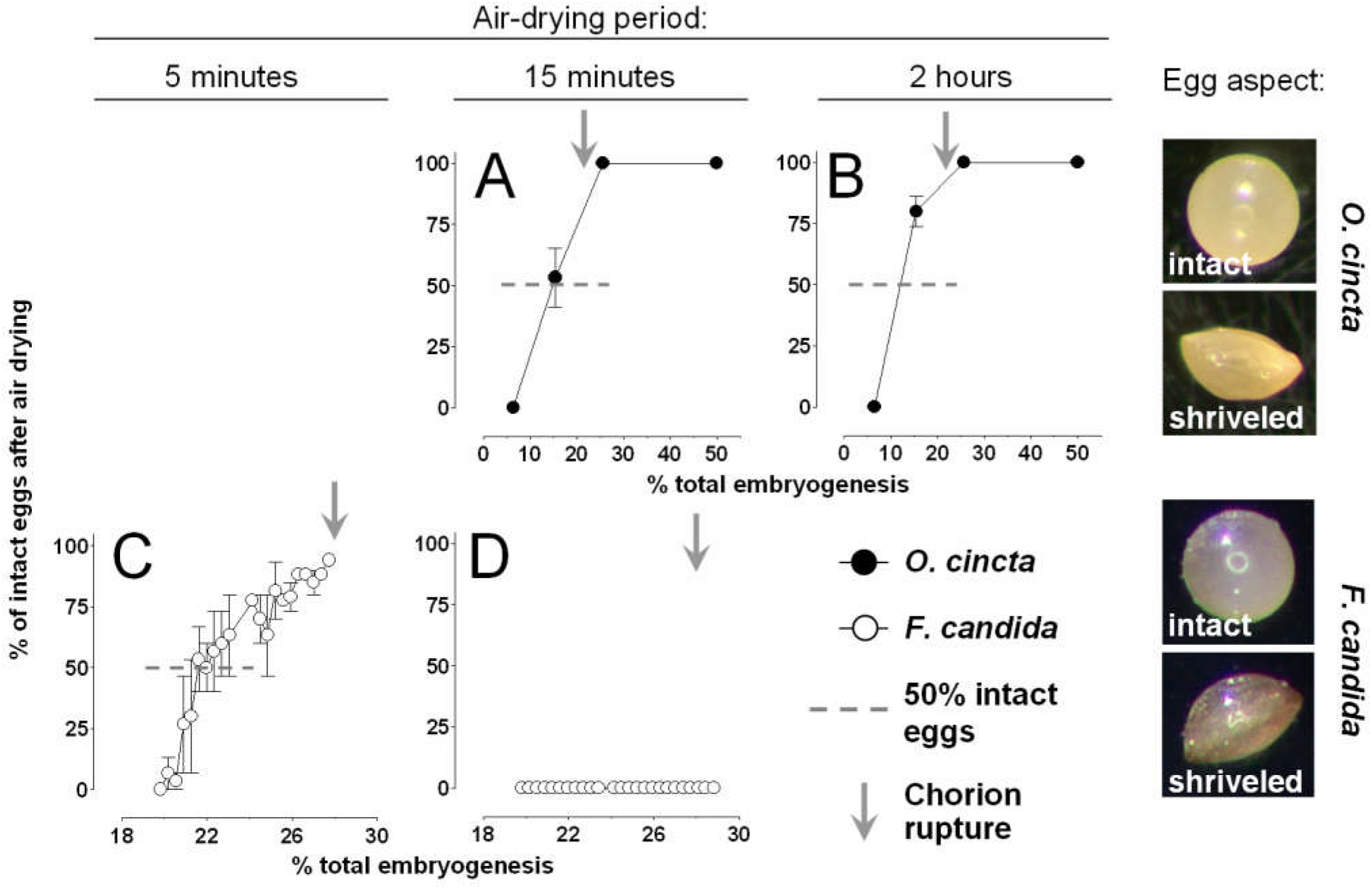
Egg permeability changes during embryogenesis at 20 ºC. (**A**, **B**) Eggs of *O. cincta* were air-dried for 15 minutes or two hours, at different embryonic ages. (**C**, **D**) Eggs of *F. candida* were air-dried for five or 15 minutes, at different embryonic ages. Air-drying periods are indicated above each panel. The percentage of intact eggs (i.e. that did not shrivel) in the horizontal axis was recorded in different samples, at distinct stages of embryogenesis indicated in the vertical axis. A total of 890 and 1435 eggs of *O. cincta* and *F. candida*, respectively, were evaluated. Grey bars indicate the period of chorion rupture (see Figure 3). Right-most panels indicate the aspect of intact and shriveled eggs.

**Table 1.**
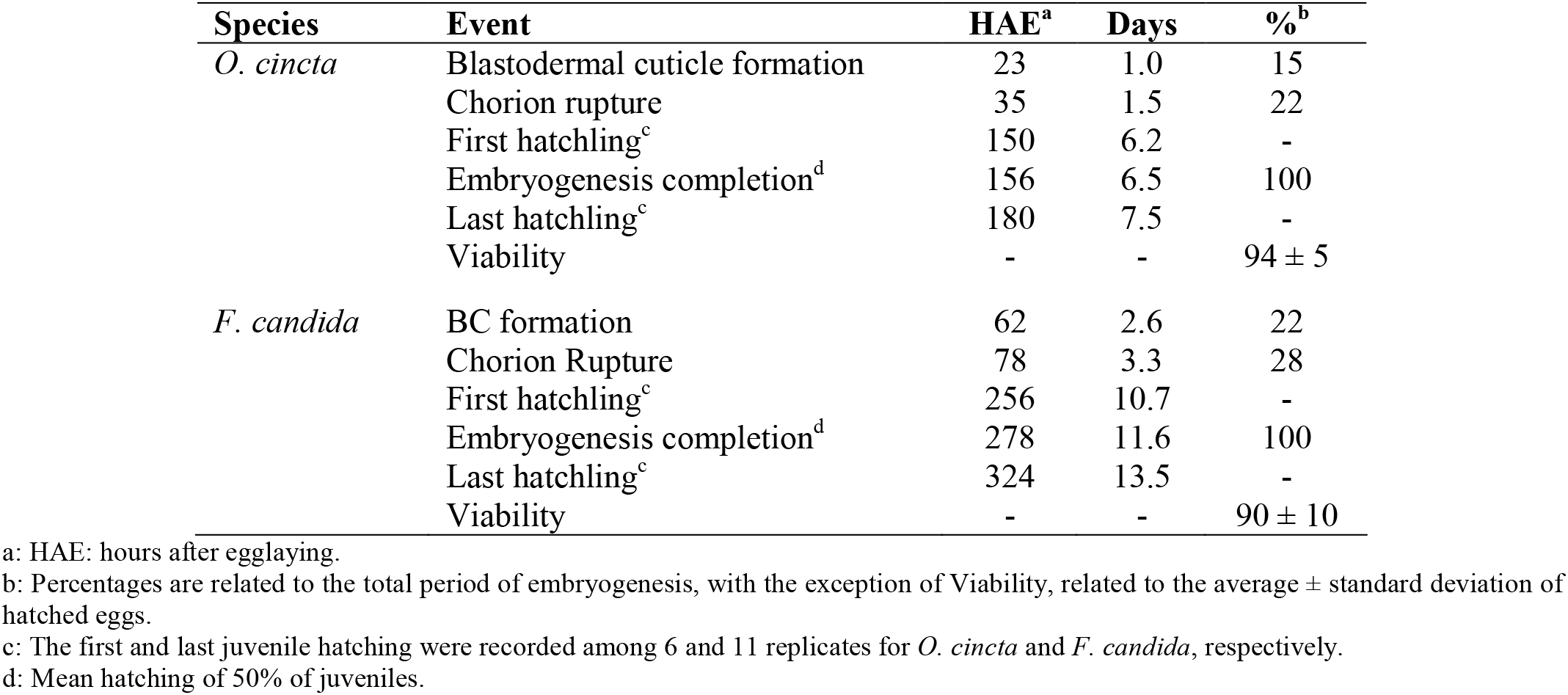
Parameters related to springtail embryonic development at 20 ºC.

### Formation of the blastodermal cuticle is related to an increase in egg viability under desiccation stress

Finally, we directly tested the relationship between blastodermal cuticle formation and egg protection against desiccation by assaying egg viability (Figure 6). Eggs developing in moist plaster were transferred at different ages to a dried filter paper to air dry for 15 minutes or 2 hours. After this period, eggs were returned to moist plaster and viability was evaluated at the end of embryogenesis in terms of hatching rate. Embryos of both species die under dry conditions before blastodermal cuticle formation, but exhibit a statistically significant increase in survival after this cuticle is formed. Eggs of *O. cincta* are comparatively more resistant and survive with high viability rates for both 15 minutes and two hours under dry conditions. On the other hand, eggs of *F. candida* are less resistant, showing high viability only for the 15-minute desiccation assay. In the two-hour desiccation experiment, eggs presented some viability only shortly after blastodermal cuticle formation (see Discussion).

**Figure 6:**
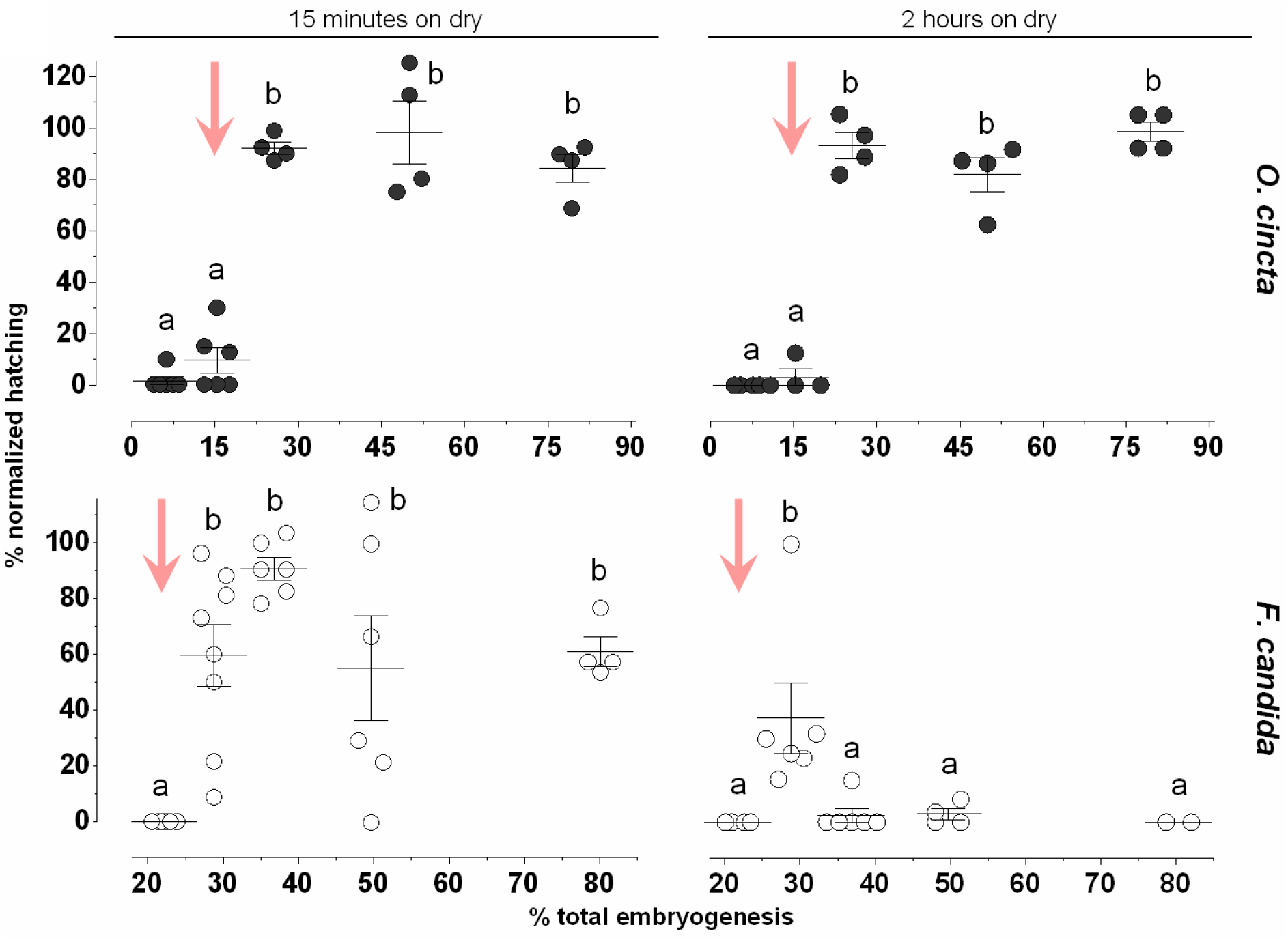
Formation of the blastodermal cuticle coincides with a protection against water loss. Eggs developing in moist at 20 ºC were transferred to dry filter paper in the moments indicated in the horizontal axis. For each time point, eggs were left drying for 15 minutes or 2 hours, when they were returned to moist until embryogenesis completion, when hatching rates were recorded. Data are expressed as percent viability, normalized from control samples, kept under moist conditions throughout development. Each circle shape represents a replicate, line and bars represent mean and standard error. Pink arrows indicate when the blastodermal cuticle is formed as determined on Figure 5. A total of 960 and 1655 eggs were employed for *O. cincta* and *F. candida*, respectively. Different letters represent significant differences among distinct stages of embryogenesis in the same drying period and for the same species, according to ANOVA followed by Tukey’s Multiple Comparison Test (P < 0.0001).

## Discussion

Our study on the potential role of the blastodermal cuticle links springtail egg developmental stages to its physiological requirements and ecological relevance. Thus, we contextualize the current findings from the perspective of environmental influences in diverse habitats and potential implications during hexapod (and insect) evolution.

Springtails are widely distributed, ranging from moist soils and tree tops to extreme habitats, such as deserts and polar regions (Greenslade, 1981; Hopkin, 1997). Two of the main abiotic factors that delimit collembolans distribution are temperature and water availability (Hopkin, 1997). There is an inverse correlation between temperature and the duration of embryogenesis (Hopkin, 1997), as observed for insect eggs (Farnesi et al., 2009). These values are known for several collembolan species, ranging from six to 80 days (Table 2) and our data are in accordance with previous descriptions for *O. cincta* and *F. candida*. Temperature also affects fecundity, egg size, resilience, and also egg dormancy. *Orchesella cincta* females that were acclimated and oviposited at 16 ºC produced eggs that are larger and more resistant to temperature variation than eggs laid from females that were acclimated and oviposited at 20 ºC (Liefting et al., 2010). *Folsomia candida* females that were acclimated and oviposited at 24 ºC produced eggs that were able to develop between 16 and 26 ºC, but not at 28 ºC. If females were acclimated at 28 ºC, a quarter of the population produced viable eggs at this temperature. Most of the females kept one month at 28 ºC did not produce eggs, but when they were returned to 24 ºC they resumed laying normal eggs after one or two days (Marshall and Kevan, 1962). Eggs of the temperate-dwelling *Isotoma viridis* are nondormant if they develop at ≥ 15 ºC but they enter a quiescent-type of dormancy, in which embryogenesis is completed but juveniles do not hatch, if they develop at ≤ 14 ºC. These quiescent eggs can hatch after 319 days if the temperature is raised to ≥ 16 ºC (Tamm, 1986). Overall, the immobile egg is a life stage that, depending on the species, lasts for weeks or months, due to a slow rate of embryogenesis or due to dormancy. During this long period the springtail egg is susceptible to predators, opportunistic microorganisms and desiccating conditions (Christiansen, 1964; Hopkin, 1997).

**Table 2.**
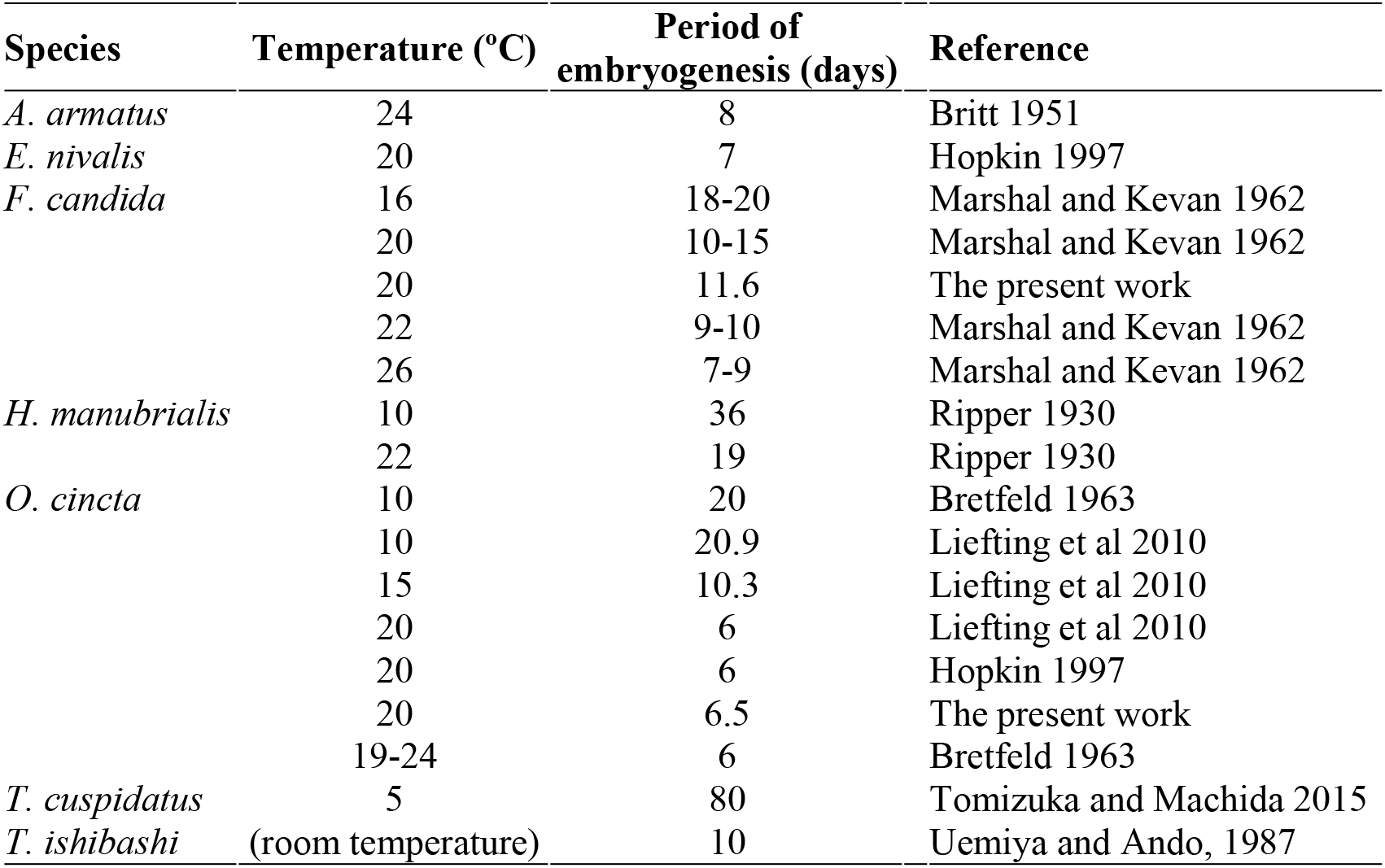
Comparative periods of springtail embryogenesis at different temperatures reported in the literature.

Desiccation resistance, *i*.*e*. the ability of an organism to survive in an arid environment, occurs through the combination of three factors: the rate of water loss, dehydration tolerance (the amount of water loss an organism can withstand prior to its death) and body water content (Hadley, 1994). Comparatively, organisms that stay alive for longer periods of drought are considered as more desiccation resistant. Due to their small size, juveniles and adults of many springtails species are prone to dehydrate when they are outside water or in an atmosphere that is not saturated with water vapor (*i*.*e*. with a relative humidity below 100%) (Hopkin, 1997; Kærsgaard et al., 2004). In these cases, different strategies can be employed to minimize desiccation stress, such as a lower rate of integument transpiration or quick water reabsorption through the ventral tube (collophore) (Figure 2F’, I) (Verhoef and Witteveen, 1980), short life cycle, anhydrobiosis or ecomorphosis (Greenslade, 1981) or water vapor absorption through the production of osmolytes (Bayley, 1999), among other mechanisms (Hopkin, 1997; Kærsgaard et al., 2004; Marx et al., 2012). In addition, the literature also describes the existence of desiccation resistant eggs, often associated with egg dormancy, as a strategy to sustain the springtail life cycle until more favorable moist conditions are available (Greenslade, 1981; Hopkin, 1997; Wallace, 1968). However, the mechanisms of how these eggs resist desiccation have not been discussed.

The first process that increases egg resistance to desiccation in springtails is water uptake (Briti, 1951; Marshall and Kevan, 1962; Uemiya and Ando, 1987a), which increases egg water content, its size and shape. For instance, *F. candida* eggs when laid have an water content of 1.12 mg / mg of dry weight and this value increases up to 2.6-fold by ~ 70% embryogenesis (Holmstrup 2019). This uptake occurs mostly during early springtail embryogenesis, prior to the formation of the blastodermal cuticle and, most likely, is essential for proper egg development, as is also the case with insects (Hinton, 1981; Rezende et al., 2016). For exemple, a similar process has been documented in insects such as crickets, a butterfly and mosquitoes (*e*.*g*. Donoughe and Extavour, 2016; Farnesi et al., 2015; Kliewer, 1961; Kobayashi, 1998; McFarlane and Kennard, 1960).

The second process that increases the desiccation resistance of springtail eggs is addressed here: a decrease in the rate of water loss that occurs during early embryogenesis (15 - 22%, Fig 5). It is understood that the blastodermal and serosal cuticles are homologous structures (Prymus-Naczyńska, 1978; Rezende et al., 2016) and it was shown that the serosal cuticle - and the chitin contained on its endocuticle - unambiguously increases egg resistance to desiccation in insects (Jacobs et al., 2013). Therefore, we interpret the present results as a direct role of the blastodermal cuticle in decreasing the rate of water loss. However, other hypotheses can not be excluded, such as the ability of the embryo *per se* and/or of the dorsal organ to have direct roles in this process (Tamarelle, 1981). In light of molecular techniques that have been recently developed for *O. cincta*, such as RNAi-driven gene silencing and *in situ* hybridization in eggs (Konopova and Akam, 2014), there is potential scope for future functional studies through targeting candidate genes, such as chitin synthase.

Meanwhile, our results show that after blastodermal cuticle formation there is a clear increase in egg viability outside the water for both *O. cincta* and *F. candida*. Despite this commonality, *O. cincta* eggs are better protected than *F. candida* ones, which is consistent with the specific microhabitats occupied by each species. *Orchesella cincta* is found in the upper levels of the litter (*i.e.*. an environment less humid than the soil) and also on the twigs of trees (Hopkin, 1997). *Folsomia candida* is found in the humus layer (*i.e.*, within the soil), a zone that usually contains 100% relative humidity (Waagner et al., 2011). In fact, when adults are compared, *O. cincta* exhibits a greater resistance to desiccation than that presented by *F. candida* (Kærsgaard et al., 2004). Curiously the water permeability (or water conductance) of the *F. candida* egg before serosal cuticle formation is six times lower than those of the adult (Kærsgaard et al., 2004, Holmstrup 2019). In addition, *F. candida* ceases to lay eggs when its surrounding relative humidity drops only slightly, from 100 to 98.8% (Waagner et al., 2011), consistent with a lack of egg viability at 98.7% relative humidity (Holmstrup, 2019). Therefore*, F. candida* adults are sensitive to water loss and their eggs present a comparatively mild desiccation resistance, necessitating a stably humid habitat throughout the life cycle.

Another trait that may decrease the rate of water loss is egg clustering: evaporation of water from eggs inside the cluster is slower than that of eggs located at the rim of the cluster. This strategy has been described for butterfly and mosquito eggs (Clark and Faeth, 1998; Clements, 1992), is also present in *F. candida* (Marshall and Kevan, 1962) (Figure 2, Suppl. Figure 3B), but absent in *O. cincta* (Figure 2).

In *F. candida* the chorion rupture occurs with 3.3 days, which is equivalent to 28% of embryogenesis (Table 1), when the embryo is still at the blastoderm stage (Figure 4B). In other words, about a quarter of total embryogenesis is dedicated to form the blastoderm and its cuticle. It is only after this cuticle has formed and expanded, providing the sole protective layer against the external environment, that the majority of embryogenesis (germ band formation and segmentation, growth of limbs and body parts and dorsal closure) proceeds. Also in *F. candida*, there is a slight decrease in egg resistance to desiccation after the blastodermal cuticle is formed (see Figure 6). This might be related to an active role of extraembryonic cells (*i*.*e*. the dorsal organ) in protecting against water loss, as previously suggested (Tamarelle, 1981). A similar function has been ascribed to serosal cells in insects (Jacobs et al., 2013), and a similar pattern is observed in the mosquito *Anopheles gambiae* (Goltsev et al. (2009); see their Figure 1B).

An interesting difference in egg protection between springtails and insects is the fact that the former is protected only by one type of layer: apart from a small period in their embryogenesis, the springtail egg is protected by only the chorion or the blastodermal cuticle. On the other hand, eggs of many insect species produce a serosal cuticle while maintaining their chorion (even for those species that uptake water and increase their egg volume - see above), thus allowing them to be protected by both types of layers at the same time. This could be relevant to allow eggs to expand their niches to more dry habitats, while most springtails species are still largely confined to humid ones. This idea that the physiological unit protecting insect eggs against water loss is the chorion in addition to the serosal cuticle was put forward over seventy years ago by Beckel (1958).

As a prospect, it will be interesting to investigate the water relations in eggs of insect species where the serosa does not produce a cuticle, such as the ecologically widespread milkweed bug *Oncopeltus fasciatus* (Dorn, 1976). Similarly, comparisons with eggs of other arthropods, such as the terrestrial crustacean *Porcellio scaber* (Whitington et al., 1993), the aquatic crustacean *Parhyale hawaiensis* (Rehm et al., 2009) and the myriapod *Trigoniulus corallinus* (Kenny et al., 2015), could elucidate the strategies employed in other lineages.

Finally, the results described here enable us to speculate on a potential terrestrialization scenario within the hexapod lineage. The soil (or other interstitial spaces) is hypothesized to be the route for hexapod land colonization (Dunlop et al., 2013; Hopkin, 1997). Consequently, a gradual process of land colonization may have occurred: the first terrestrial hexapods were very desiccation sensitive and lived in moist soils. Those that acquired traits to prevent water loss could go above the soil and live in the litter or other epidaphic environments. From there, the groups that acquired further traits to withstand even harsher desiccating conditions could leave the epidaphic environment and conquer drier niches, even xeric ones, such as deserts. Whether *F. candida* has always remained in moist soils or it returned to this niche subsequently during its evolutionary history is not yet possible to evaluate. In addition, blastodermal cuticle might also protect springtail eggs against excessive water uptake during flooding, when eggs of some species are kept submerged for long periods in a dormant state (Tamm, 1986, 1984).

## Concluding remarks

The main role of the eggshell is to protect the developing embryo. During Tetrapoda evolution, for example, a calcified eggshell was essential to minimize water loss facilitating evolution on land (Dunlop et al., 2013). We consider that convergent solutions also arose in the hexapods. The role of the blastodermal cuticle in springtail egg resistance to desiccation would be similar to the function undertaken by the serosal cuticle in insect eggs. Given that the blastodermal cuticle is present in species within all the other arthropod groups (Rezende et al., 2016) we envisage that the production of this structure in aquatic ancestors was an important prerequisite for the evolutionary process of Hexapoda terrestrialization. After the split of springtail and insect lineages, insect evolution led to the acquisition of two extraembryonic epithelia (amnion and serosa), an amniotic cavity and, often, a serosal cuticle (Machida and Ando, 1998; Panfilio, 2008; Rezende et al., 2016). As previously suggested (Jacobs et al., 2013; Zeh et al., 1989), the presence of these traits as well as a chorion that does not rupture equipped insects with the possibility to exploit new niches and attain an adaptive success that has no parallel among other arthropods and even other animals.

## Acknowledgments

This study was financed by the Coordenação de Aperfeiçoamento de Pessoal de Nível Superior - Brasil (CAPES) - Finance Code 001 and FAPERJ. H.C.M.V. was supported by a fellowship from CAPES (88881.132450/2016-01). The authors are also indebted to Janine Mariën for technical assistance. G.L.R. thanks Ryuichiro Machida (Sugadaira Research Station, Mountain Science Center, University of Tsukuba) for providing access to the 1963 Bretfeld paper. We also thank Ademir de Jesus Martins Júnior (IOC, Fiocruz), Clicia Grativol (CBB, UENF) and Gabriel Costa Queiroz (Museu Nacional, UFRJ) for critical reading and suggestions on the manuscript. K.A.P. thanks Michael Akam (Department of Zoology, University of Cambridge, UK) for access to a laboratory culture and resources in support of the descriptive *Folsomia candida* embryology work.

## Supplementary Figures

**Supplementary Figure 1.**
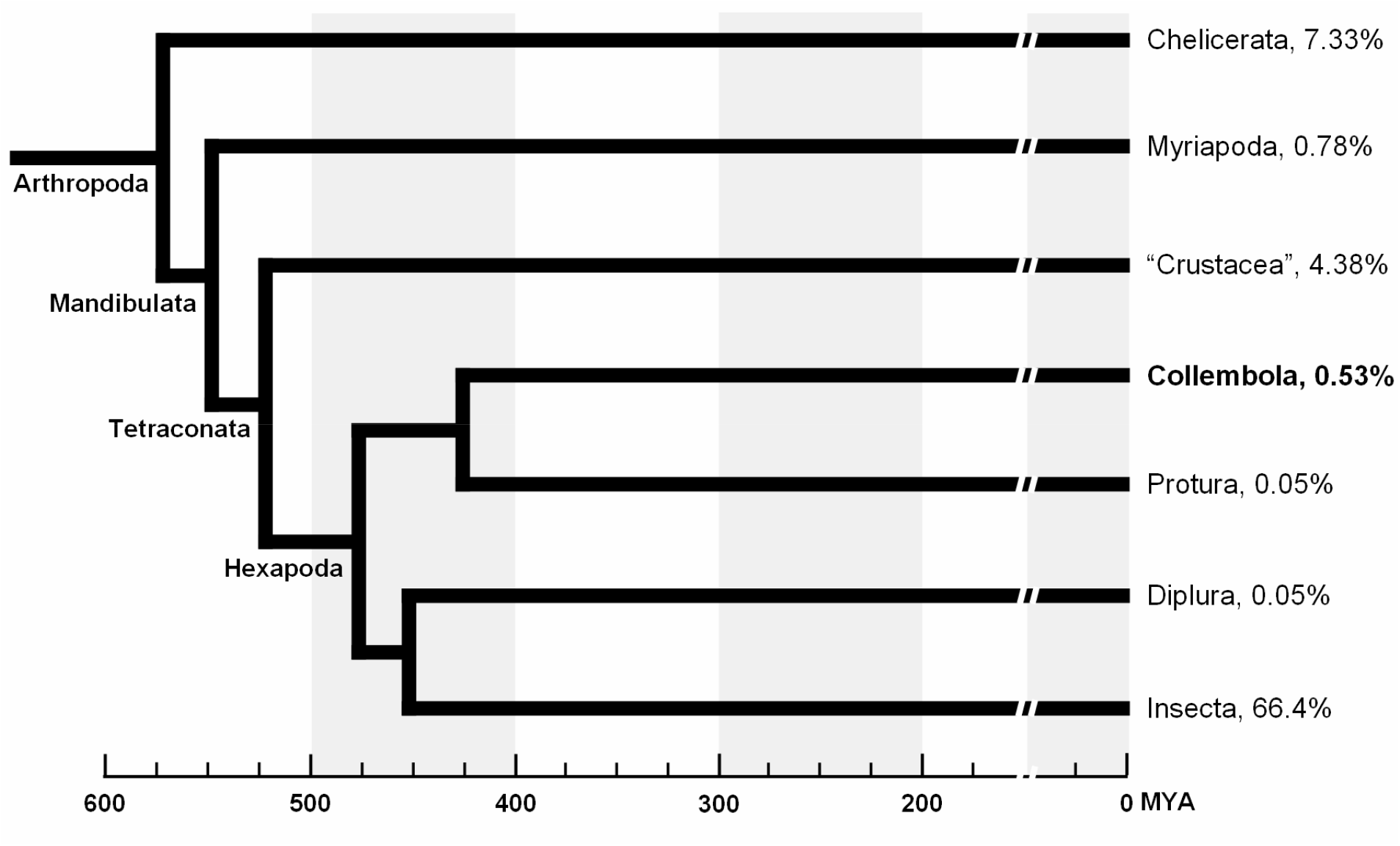
Arthropod phylogeny with estimates on time of divergence and number of species. The divergence periods indicated are the median time estimated at timetree.org (accessed on 27/03/2019). Values alongside each terminal taxon indicates the percentage of species regarding the total number of living animal species, estimated as 1,527,660 (Zhang 2011).

**Supplementary Figure 2:**
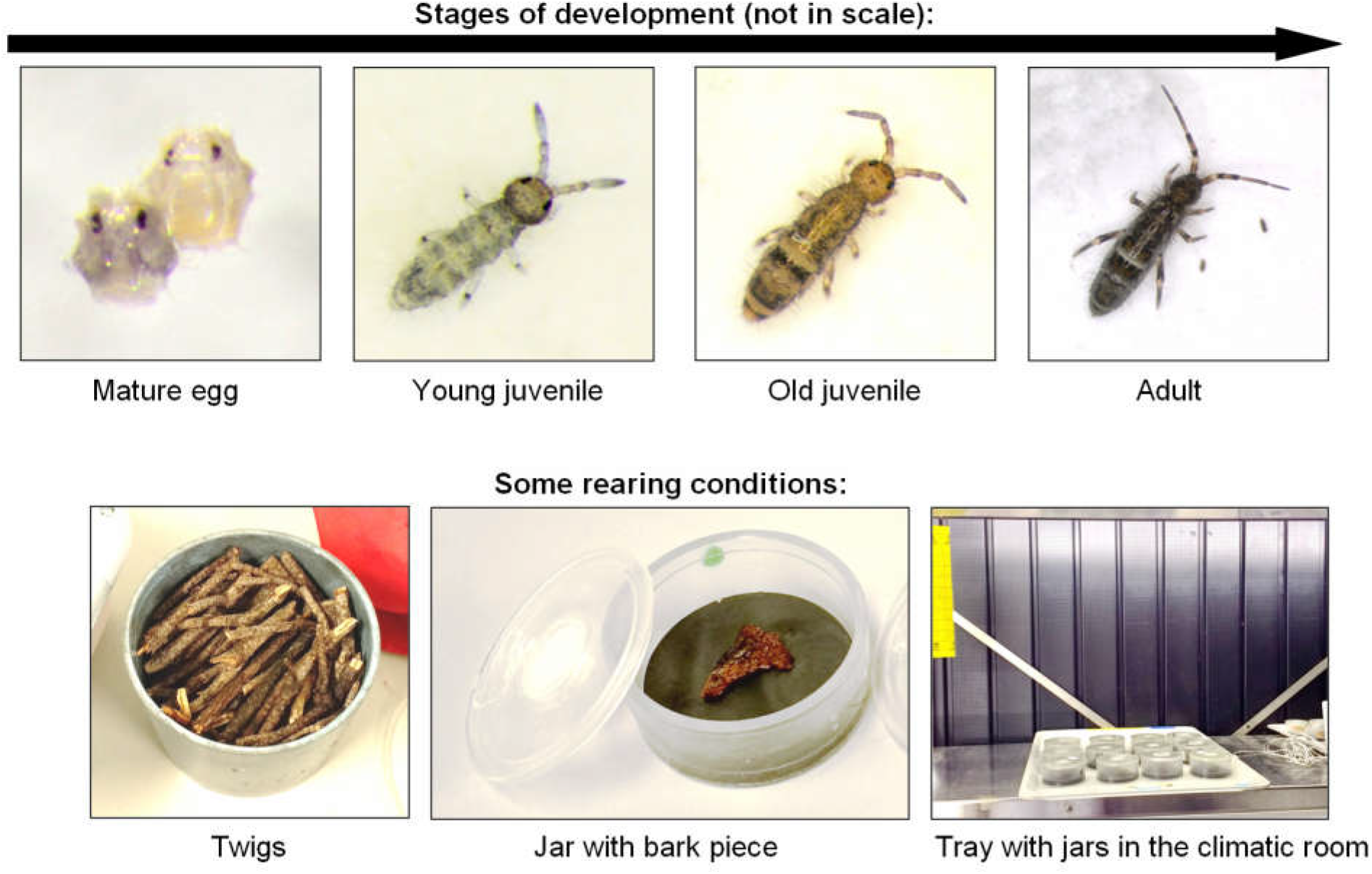
Background information for *Orchesella cincta*.

**Supplementary Figure 3.**
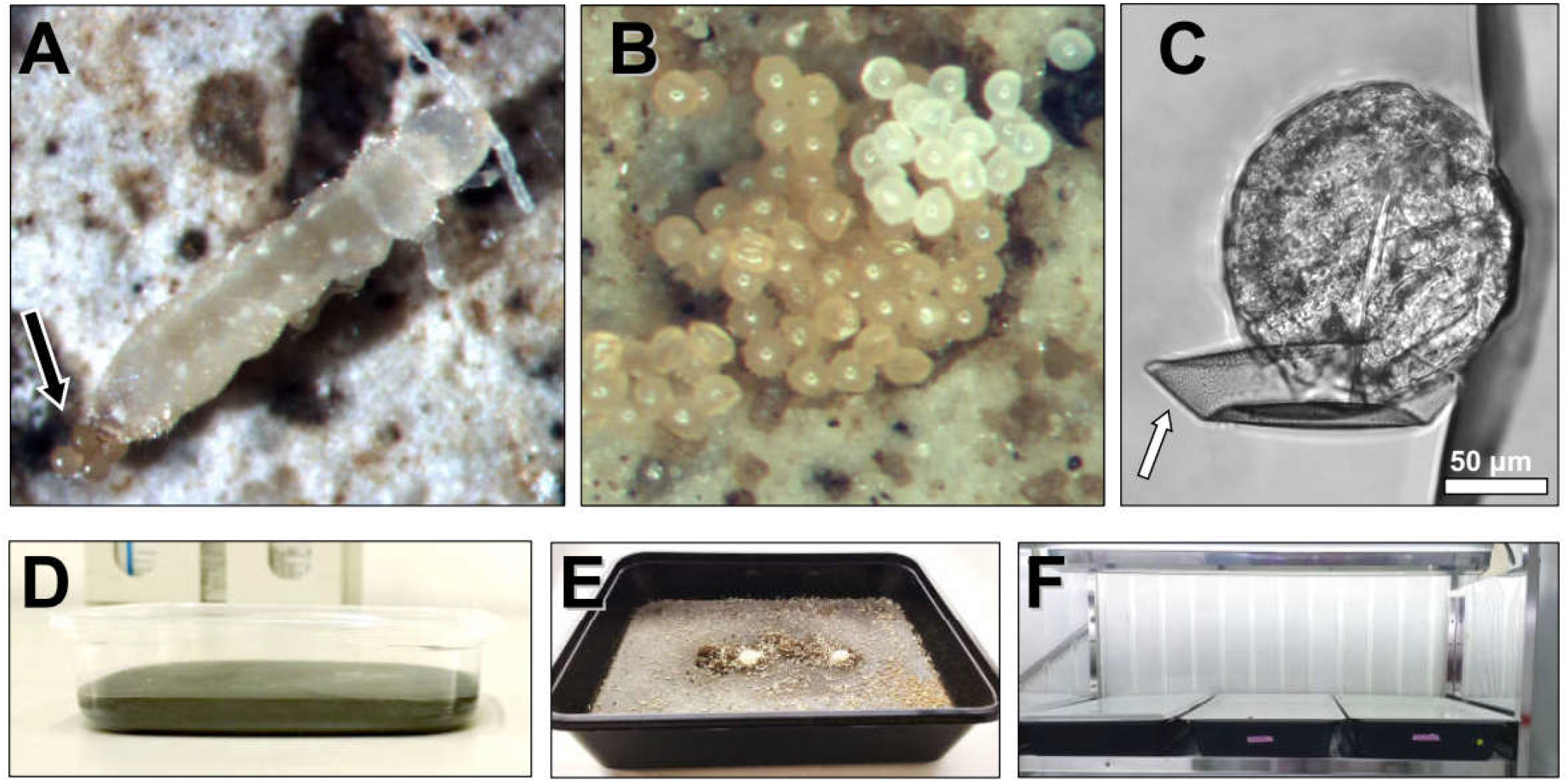
Background information for *Folsomia candida*. (**A**) Parthenogenic female laying eggs (arrow). (**B**) Cluster of eggs: freshly laid eggs are white and they become tan-colored (*i.e.* yellowish) after a while. (**C**) DIC microscopy image of an egg after chorion rupture. Arrow indicates a polar cap (see also main Figures 1 and 3). The upper polar cap is missing. (**D**) Jar with plaster of Paris and charcoal. (**E**) Large jar with adults (**F**) Jars in the climatic room.

**Supplementary Figure 4.**
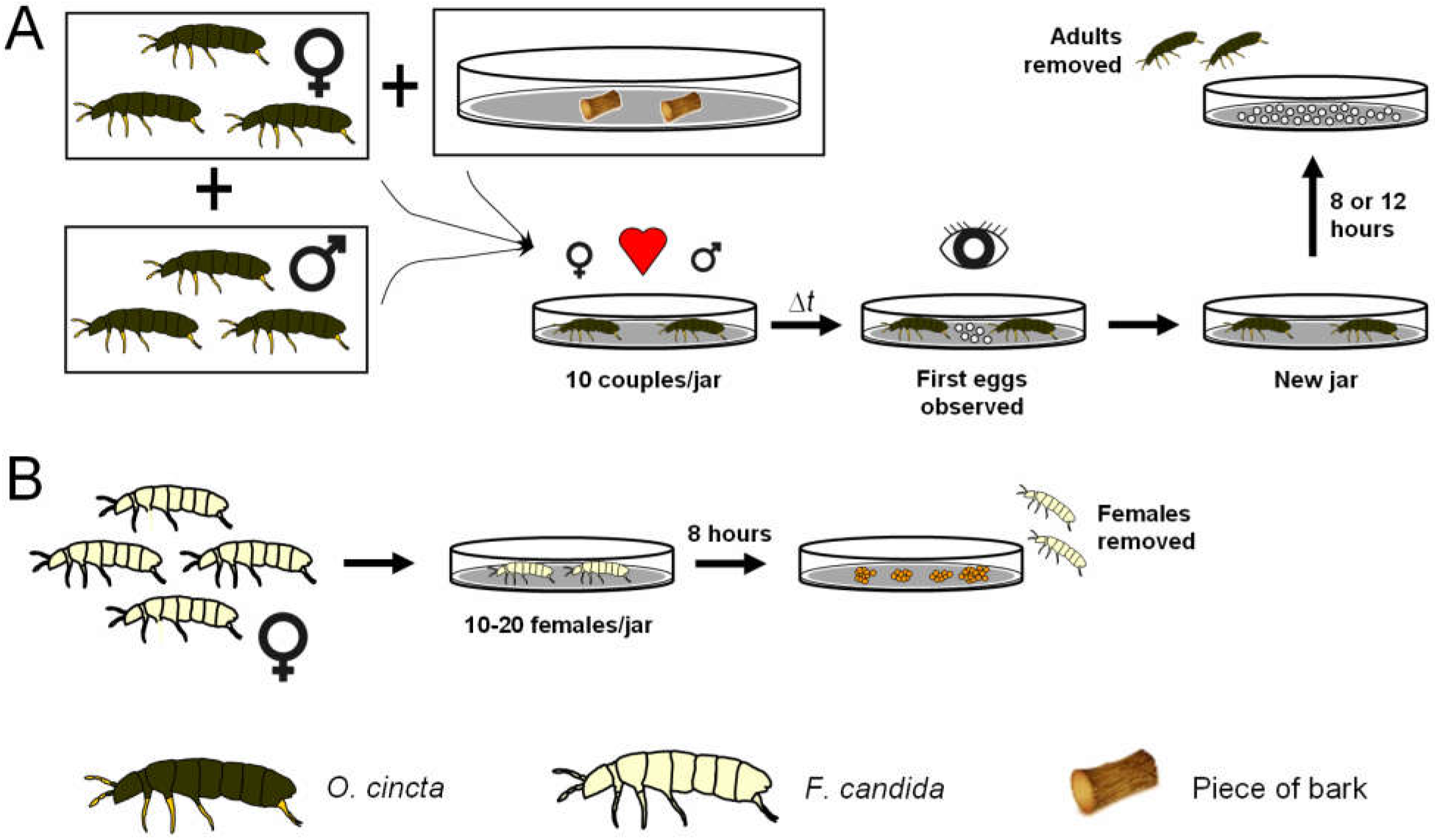
Scheme of *O. cincta* and *F. candida* egg laying. (**A**) *O. cincta* adults were sexed and transferred to small jars containing a piece of bark. Jars were carefully observed to check the presence of the first eggs. Adults were then transferred to a new jar where egglaying lasted for 8 or 12 hours, when adults were removed. (**B**) Parthenogenic *F. candida* females were transferred for small jars, where egg laying lasted for 8 hours, when females were removed.

**Supplementary Figure 5.**
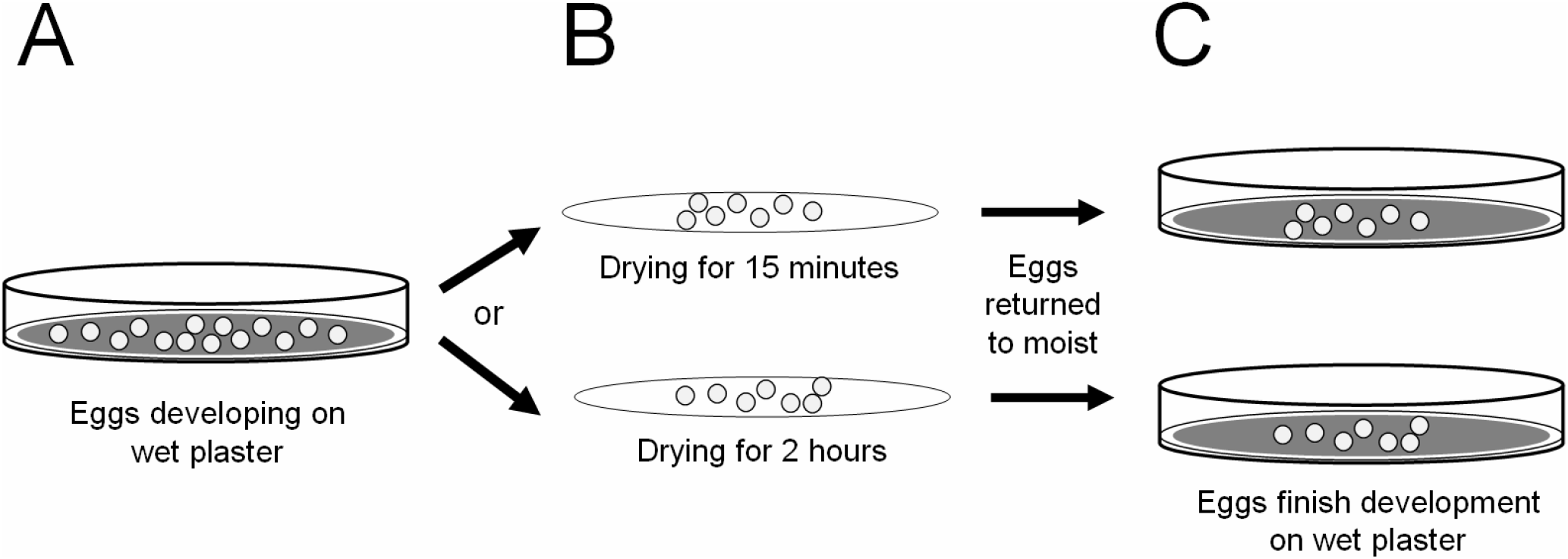
Methodology for the egg viability assay. The procedure was the same for *O. cincta* and *F. candida*. (**A**) Eggs are developing on wet plaster until the desired stage is reached. (**B**) Eggs were then left air-drying for two different periods: 15 minutes or 2 hours. (**C**) After these periods were completed, eggs were returned to wet plaster until the completion of embryogenesis, when hatching rates were scored. Controls were left on wet plaster throughout the whole embryogenesis (not shown).

